# Early amyloid spine response and impaired synaptic transmission of pyramidal neurons in human biopsies with Alzheimer’s Disease-related pathology

**DOI:** 10.1101/2025.01.07.630516

**Authors:** Antonios Dougalis, Polina Abushik, Anssi Pelkonen, Luca Giudice, Mireia Gómez-Budia, Nataliia Novosolova, Nelli-Noora Välimäki, Mohammad Rezaie, Dilyara Nurkhametova, Raisa Giniatullina, Anastasia Shakirzyanova, Akash Mali, Tuomas Rauramaa, Beth Stevens, Mikko Hiltunen, Ville Leinonen, Tarja Malm

**Author notes:** Equal contribution (Antonios Dougalis, Polina Abushik).

## Abstract

Studies of neuronal functions during the pathological progression of Alzheimer’s disease (AD) in humans are limited due to the lack of live human brain tissue from patients with AD. To address this gap, we have established an exceptional approach to study the electrophysiological properties and cell morphologies of human neurons in acute slices obtained from cortical biopsies of patients with idiopathic normal pressure hydrocephalus (iNPH). Histological examination of Broadman area 8-9 cortical biopsies from these patients have revealed that approximately 40% of the patients show signs of early AD-related pathology in the form of low to moderate, often fleecy beta-amyloid (Aβ) deposits and additional, occasional tau in 10% of the cases. Thus, the iNPH brain biopsies, obtained during the shunt surgery to treat the patients, offer a unique window to investigate how existing AD-related pathology alters the operational properties of human cortical neurons. Here we carried out integrative analysis of human neuronal electrophysiology at single neuron and network level followed by subsequent cellular morphological reconstructions to register the primary pathological changes in neuronal functions in correlation with existing AD-related pathology. The presence of Aβ plaques induced a decrease in basal excitatory synaptic activity in pyramidal neurons residing on supragranular layers of the cortex. These neurons received less of L1-induced inhibition and appeared hyperexcitable in response to application to NMDA in multielectrode array (MEA) recordings. Interestingly, the global spine density of supraganular pyramidal neurons was increased in biopsies with AD-related pathology. The increase in spine density was coincidental with a partial recovery of excitatory transmission (frequency but not amplitude), of L1-induced inhibition in supragranular layers pyramidal neurons and of NMDA induced supragranular firing (but not of bursting hyperexcitability) indicating a potential differential effect of tau in the presence of Aβ on the progression of neuronal functions. Despite the partial renormalization of deficits seen in cases with Aβ pathology only, pyramidal neurons in cases with both Aβ and tau exhibited more consistent deficits in the intrinsic neuronal properties with increase in sodium and potassium currents and a strong propensity to bursting under NMDA stimulation. We conclude that complex mechanisms operate in response to accumulation of Aβ and tau including re-structuring of the apparatus of synaptic transmission and consolidation of a hyperexcitable supragranular cortical network phenotype. The observed changes in spine density and synaptic activity are reminiscent of parallels seen in homeostatic plasticity and synaptic scaling and may depend on strong interactions with the local microenvironment (astrocytes and microglia). This is the first study to report the impact of AD-related pathology on single-neuron operational properties and morphology in humans.

## Introduction

Despite extensive research, the mechanisms conferring to Alzheimer’s disease (AD) progression in humans are still unclear. Studies of the pathological processes in the human AD brain are very limited due to the lack and inaccessibility of live human brain tissue from patients with AD. Here, we studied in detail the electrophysiological and morphological properties of live human neurons in acute slices obtained from cortical biopsies from patients with idiopathic normal pressure hydrocephalus (iNPH).

iNPH is a condition caused by the inability of cerebral spinal fluid (CSF) to circulate unobstructed in the ventricular space. This leads to the classical clinical triad symptoms in iNPH patients, postural instability, incontinence and cognitive deficits^1,2^. iNPH is treated by shunt surgery to bypass the accumulating CSF from the lateral ventricle into the abdomen^3^. During the surgery, a brain mini biopsy (10-20 mm^3^) was excised for diagnostic purposes at the shunt insertion site. iNPH and AD are intertwined so that half of the iNPH patients show signs of early AD-related pathology in their surgical brain samples, evidenced as fleecy Aβ deposits in 40% of the patients and additional, occasional tau in 10% of the cases ^4,5^. Many of these patients eventually develop clinical AD in the next 1-5 years^6–8^. Extensive literature confirms that iNPH patients show the natural course of AD progression both in pathology and clinical disease^*6–10*^. Due to the early AD pathology present in half of the iNPH patients, iNPH patient brain biopsies offer a unique window to evaluate events occurring during the development of early AD.

Single cell RNA sequencing datasets on human post-mortem AD brains and rodent models of AD have revealed that brain cells exist in disease-specific cellular states ^11–13^ that are fundamentally different between mice and humans^14–16^. However, post-mortem human tissue suffers from post-mortem artifacts^17^, represents only the final stages of the disease and does not allow any functional physiological studies since the tissue cannot be preserved alive.

Traditional animal AD models are crucial in research but have limitations that prevent them from fully capturing the complexity of human neurodegeneration^18^. For instance, rodent models often depend on genetic modifications to replicate amyloid or tau pathologies, while they do not reflect the intricate interactions found in human neural microenvironments, making it challenging to reconcile and apply findings directly to human AD, particularly in terms of understanding early-stage cellular and molecular dynamics^18^. In contrast, the biopsy specimen from iNPH patients provide a meaningful translational model that preserves native human-specific tissue architecture, cellular diversity, and gene expression patterns, leading to insights that are unattainable through animal studies.

Recent research indicates that Alzheimer’s pathology is associated with significant declines in synaptic formation and function, attributed in part to dysregulation within microglial populations^19^. Progressive AD is characterized by an early and late response stage in cell specific molecular signatures^20,21^ Our study here aimed to identify prominent functional neurophysiological and neuromorphological signatures in live cortical neurons in the presence of AD-related pathology in order to gain insight on the functional changes in the cortical network induced by AD-related pathology.

## Results

### AD-related pathology alters the stable intrinsic electrophysiological properties of L2/3 pyramidal neurons in iNPH patient frontal cortex biopsies

We started out assessment of neuronal function by comparing the intrinsic passive and active electrophysiological properties (Fig.1) of human pyramidal neurons in L2/3 cortical layer in iNPH patients with only Ab pathology, both Ab and tau pathology, and in patients with no AD-related pathology.

**Fig. 1.**
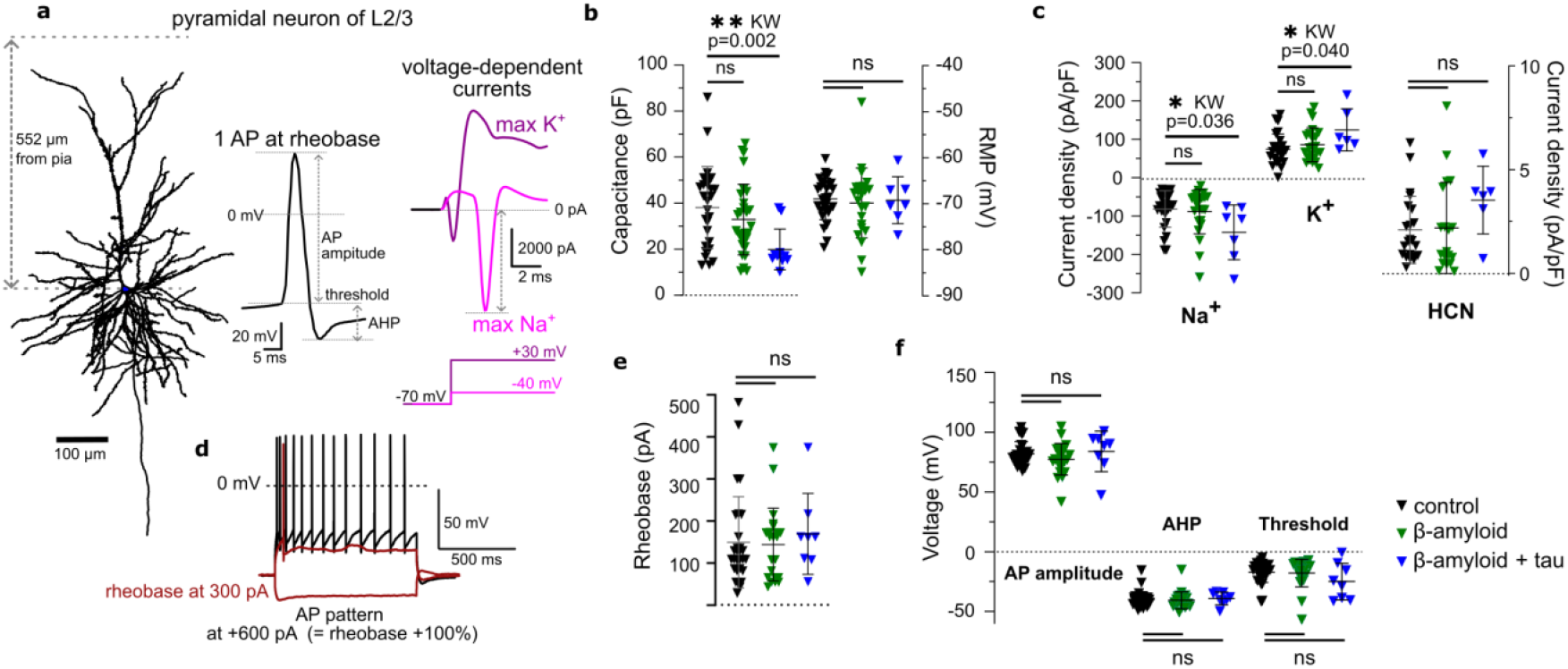
Comparative electrophysiological properties of L2/3 pyramidal neurons in iNPH patient frontal cortex biopsies. **a**. Neuromorphological reconstruction of a representative pyramidal neuron (PN) located in the L2/3 cortical layers (552 µm depth from PIA) obtained from an iNPH patient’s frontal cortical biopsy with an AP, produced by this cell and voltage-dependent currents of this cell to evaluate Na^+^ and K^+^ currents. **b**. The capacitance of pyramidal neurons of L2/3 and their resting membrane potential (RMP) according to Aβ presence in a tissue. **c**. The current density of Na+, K+, and hyperpolarization-activated cyclic nucleotide-gated cation (HCN) current in pyramidal cells registered in iNPH samples without Aβ (black), with Aβ (green), or with Aβ plus tau peptide (blue). For b, c data presented as mean±SD with statistical comparison by Kruskal-Wallis test with Dunn’s correction, ns>0.05, *p<0.05, n=26, 23 and 7 pyramidal cells/group, taken from control negative 17 biopsies, Aβ 15 biopsies and Aβ with tau 12 biopsies. **d**. AP of the pyramidal cell at the rheobase (red) and AP firing pattern (black) induced by application of current, corresponding to +100% to rheobase. **e**. The level of rheobase for PN. **f**. The maximal amplitude of AP (Max AP), afterhyperpolarization (AHP), and threshold of AP in PN. For e, f, g data presented as mean±SD with statistical comparison by Kruskal-Wallis test with Dunn’s correction, ns>0.05, *p<0.05, **p<0.01, ***p<0.001, ****p<0.0001 n=26, 20 and 8 pyramidal cells/group.

Analyses of passive membrane potential responses to small current input revealed that the whole cell capacitance (Cw) of pyramidal neurons in brain biopsies is decreased in cases with both Ab and tau (Fig.1 b), implying that the effective somatic surface area of the pyramidal neurons is shrinking as Cw is related to cell body surface ^22,23^ (see also, Extended Data Fig. 2 a, b). This change was not accompanied by an effect on the resting membrane potential with the average value remaining at about -70 mV (Fig. 1 b).

We assessed the activation of voltage-gated Na^+^and K^+^channels responsible for the repetitive APs^24^ firing by applying appropriate depolarizing voltage step protocols to evaluate their maximal current response. The maximal Na+ current appeared during membrane depolarization from -40 mV up to 0 mV, the maximal sustained K^+^ current was taken from the highest depolarization at 30 mV^24^ (Fig. 1 a, Extended Data Fig. 1 a). The analysis of voltage input to pyramidal neurons in L2/3 revealed increases in Na+ and K+ current densities in biopsies with Aβ and tau compared to biopsies containing Aβ only or no-pathology containing control samples (Fig. 1 c). We also evaluated the impact of the pathology on hyperpolarization-activated cyclic nucleotide-gated cation (HCN) channels which are known to play a role in cell excitability at the soma^25–27^ and sculpturing synaptic transmission. Analysis of HCN current as a maximal somatic current amplitude during membrane hyperpolarization from -90 to -140 mV did not reveal significant changes in HCN current density between patient groups (Fig. 1 c, Extended Data Fig. 1 b).

AP firing rate was analyzed in response to short (1 s) current inputs (input -output relationships). The rheobase, the minimal current required for a neuron to fire, was determined for every pyramidal neuron recorded in the supragranular layers (Fig.1 d, e, Extended Data Fig. 1 c, d). There were no differences in the rheobase between the different patient groups (Fig.1 d, e). We used every pyramidal neuron’s rheobase for step current normalization and performed a current load survey study where injected current pulses were delivered with current intensity of 2x, 3x and 4x the intensity of the rheobase pulse (Extended Data Fig. 1 d, e). Pyramidal neurons of L2/3 from biopsies containing Aβ only could maintain an adequate input gain response of action potential firing (Extended Data Fig. 1 e) to similar level as those from no-pathology containing control samples. Measurements of the characteristic properties of APs, such as amplitude, threshold, and afterhyperpolarization (AHP) did not reveal any significant changes between the control group and biopsies containing Aβ only (Fig. 1 f, Extended Data Fig. 1 c). The results indicate that when only Aβ pathology is present, the supragranular pyramidal neurons are characterized by relatively stable membrane properties and input output gains when compared to control neurons. However, the passive (capacitance) as well active membrane properties (sodium and potassium currents) are both disturbed in presence of concomitant Aβ and tau pathology.

### L1 mediated inhibition in supragranular layers is impaired in cases with Aβ pathology

Based on our results published in Gazestani and colleagues^20^, loss of L1 residing, NDNF1 interneuron population is an important early event in AD. To assess neuronal network inhibition stemming from L1 in cortical biopsies, we performed a stimulation study (Fig. 2 a, b) in which pyramidal neurons were current clamped to positive potentials with the help of injection of small currents to evoke a steady stream of APs (Fig. 2 c). The L1 cortical layer was stimulated to elicit an inhibitory postsynaptic potential (eIPSP, Fig. 2 d) while pyramidal neurons in L2/3 were firing under a depolarizing current. Inhibition was assessed as the mean change in pyramidal neuron firing rate elicited after three successive L1 stimuli.

**Fig. 2.**
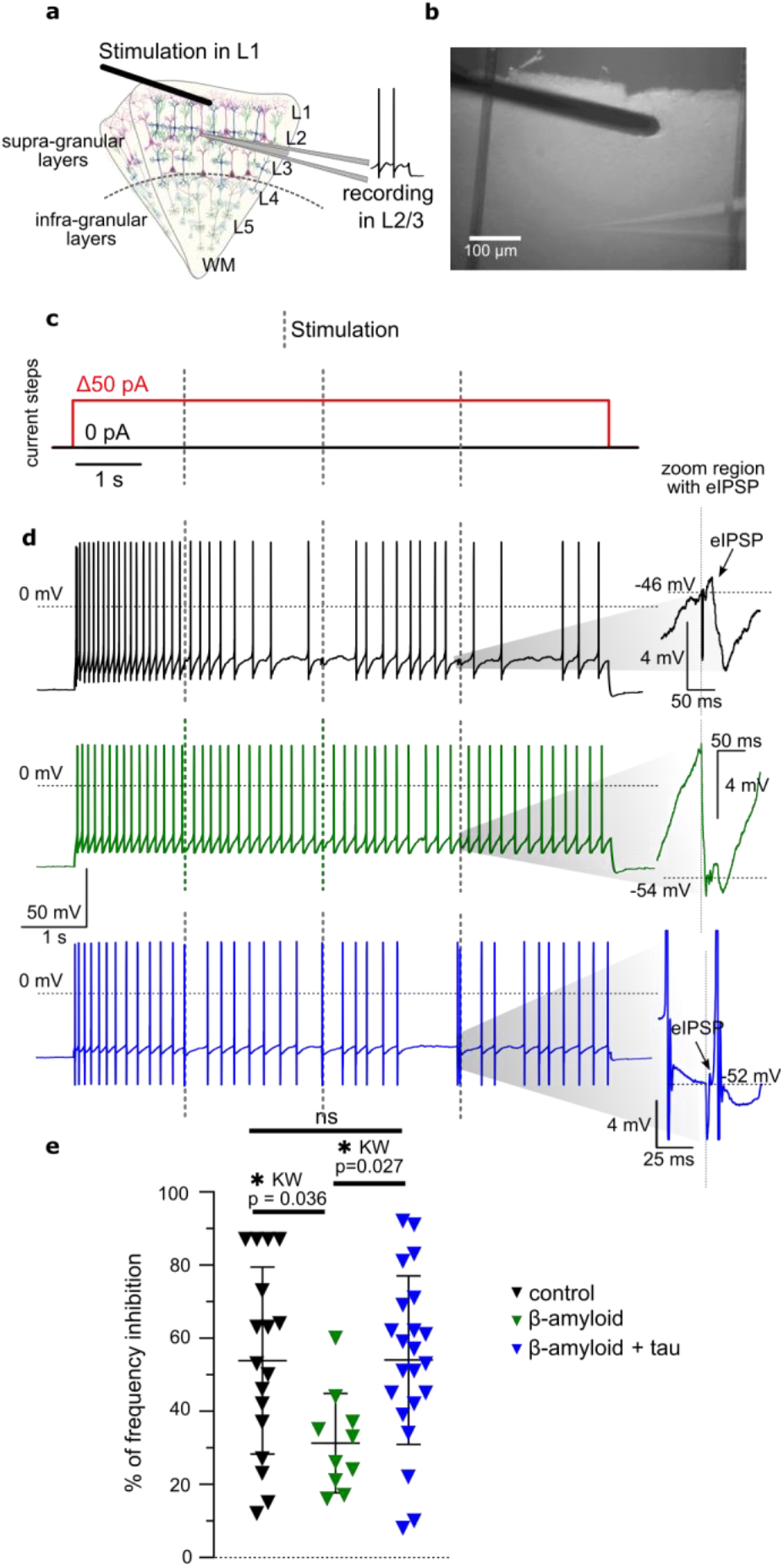
Inhibition of L2/3 pyramidal neuron activity by stimulation of L1 layer. **a**. Layer distribution scheme and image (**b**) of stimulation electrode position in L1 layer (black) and recording electrode placed in L2/3 layer in an acute slice of iNPH tissue. **c**. Protocol for current-clamp mode recording of potential during current step +50 pA (red) and 0 pA (black). Stimulation impulses are marked with a grey dotted line. **d**. Representative AP firing of PN to the current application and L1 stimulation registered in iNPH samples without Aβ (black), with Aβ (green), and with Aβ plus tau peptide (blue). Stimulation was on 1,6 s of the current step, then repeated twice at a rate of once every 2 s. Left side: zoomed regions with inhibitory postsynaptic potentials evoked by stimulation of L1 (eIPSP, marked with black narrows) **e**. The plot presents % of frequency inhibition in PN after the stimulation pulse in the L1 layer in iNPH samples without Aβ (black), with Aβ (green), or with Aβ plus tau peptide (blue). Data presented as mean±SD with statistical comparison by Kruskal-Wallis test with Dunn’s correction, *p<0.05, n=17, 10 and 22 pyramidal cells/group, taken from control negative 10 biopsies, Aβ 5 biopsies and Aβ with tau 9 biopsies.

In control biopsies L1 induced inhibition of AP firing rate was effective and amounted to about 55% (Fig. 2 d, e), while the inhibition in biopsies with Aβ only was very low, just over 30% (Fig. 2 d, e). In biopsies with both Aβ and tau, inhibition of AP firing rate was at the level of control values (Fig. 2 d, e). The lack of L1 inhibition in samples containing Aβ but not tau confirms the reported loss of L1 NDNF interneurons described previously^20^ and lend functional evidence over the disinhibitory effect of L1^21^ contributing to the hyperexcitability of the supragranular layer.

### AD-related pathology is associated with an increase in global spine density of pyramidal neurons in L2/3

Previously patch-clamped L2/3 pyramidal neurons, filled with biocytin during recordings, were used for post-hoc morphological reconstructions using Neurolucida software (Fig. 3 a-c). The spine density of pyramidal neurons in the L2/3 cortical layer was significantly increased in biopsies with AD-related pathology (Fig. 3 d, 0.105±0.062 spine/μm in control, 0.174±0.073 spine/μm in Aβ only and 0.188±0.117 spine/μm in Aβ with tau samples with comparison by Kruskal-Wallis test with FDR correction where *p<0.05 & q<0.05, n=14, 10 and 9 pyramidal cells/group, taken from control negative 12 biopsies, Aβ positive 9 biopsies and Aβ with tau positive 6 biopsies). The biggest contribution to the increase came from the increase in spine density on the apical dendrite, the main receiving process that typically intersects several of the upper layers^28^ (Fig. 3 e, Extended Data Fig. 2 d). Moreover, the analysis of the total number of spines and the length of the dendritic tree confirmed that the spine density increase was mediated to a greater extent by an increase in their number rather than by a change in the dendric length (Fig. 3 e, Extended Data Fig. 2 c, d).

**Fig. 3.**
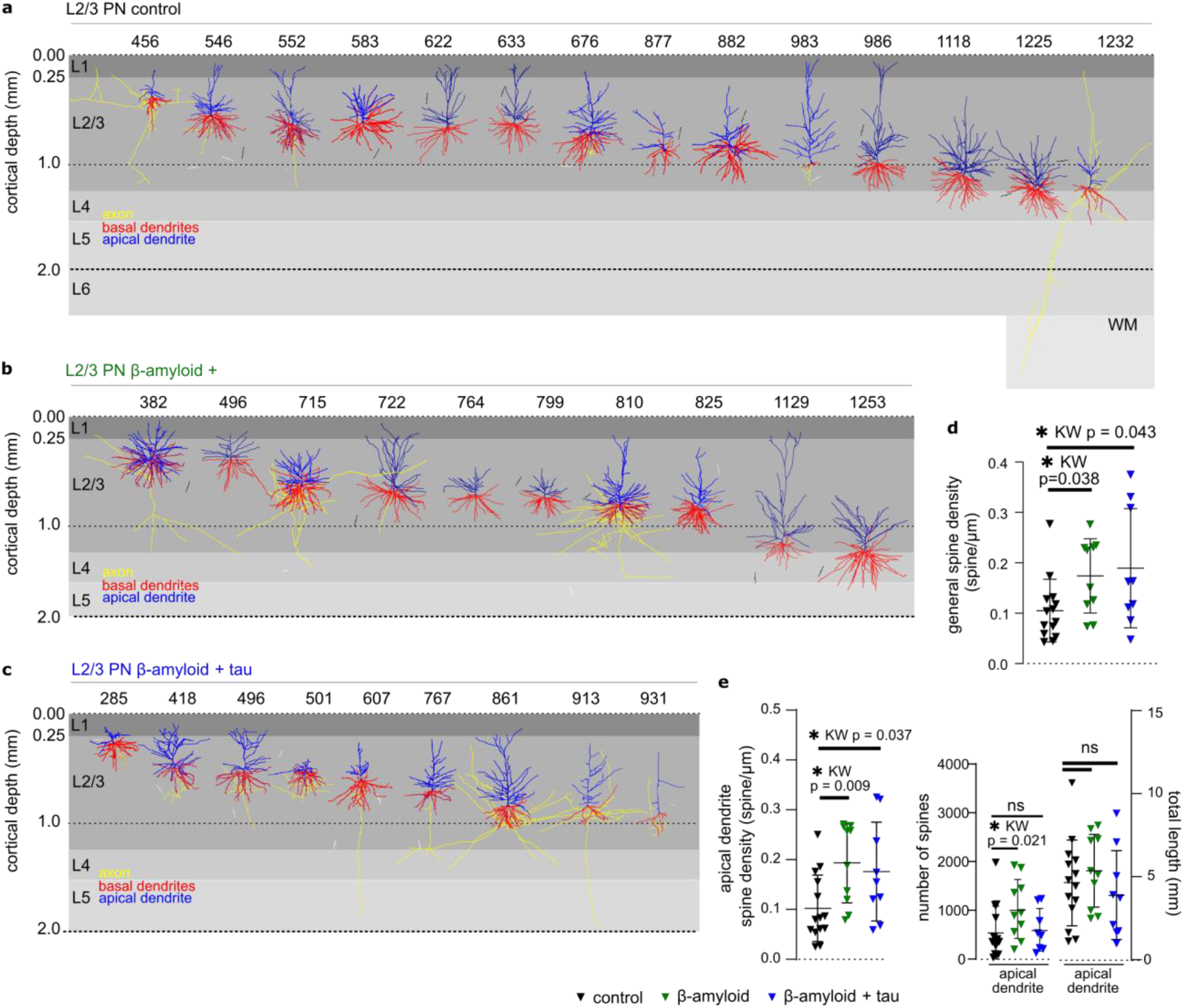
Morphology reconstruction and spine density analysis of L2/3pyramidal neurons (L2/3 PN) in iNPH patient frontal cortex biopsies. **a**. Morphologically reconstructed, biocytin-filled L2/3 pyramidal neurons from frontal cortex biopsies of control negative biopsies, (**b**) Aβ positive biopsies and (**c**) Aβ with tau positive biopsies. Somatic depth is depicted on the top bar while apical and basal dendrites of pyramidal cells are colored blue and red respectively for visualization, with axons colored yellow. **d**. L2/3 pyramidal neurons general spine density in connection with Aβ presence in a tissue. **e**. Spine density of L2/3 pyramidal neurons in the apical dendrite, the total number of spines on apical dendrites, and length of apical dendrites of L2/3 pyramidal neurons registered in iNPH samples without Aβ (black), with Aβ (green) and with Aβ plus tau peptide (blue). Data presented as mean±SD with statistical comparison by Kruskal-Wallis test with FDR correction, *p<0.05 & q<0.05, n=14, 10 and 9 pyramidal cells/group, taken from control negative 12 biopsies, Aβ positive 9 biopsies and Aβ with tau positive 6 biopsies.

Since Aβ is central to AD and part of iNPH biopsies contain Aβ or Aβ and tau, it was interesting to study how Aβ plaque might affect spine density in the immediate vicinity of the plaques. For this analysis, slices containing Aβ and reconstructed neurons filled with biocytin were secondarily stained with antibodies to Aβ (WO-2, Extended Data Fig. 3 a-d). We found a non-significant tendency towards a reduction in the local spine density in a region in close contact with Aβ (Extended Data Fig. 3 e-g). No such reduction in spine density was observed in biopsies containing Aβ and tau (Extended Data Fig. 3 g), ^29,30^ due to high variability and low number of cases available for such analysis.

### AD-related pathology suppresses the spontaneous synaptic activity of pyramidal neurons in L2/3

Since we identified an increase in spine density in early AD, it was important to clarify the functionality of these spines. To do so, we employed patch clamp recordings to identify basal excitatory and inhibitory activity as described previously^29^. It would be expected that a greater number of spines should result in an increase in spontaneous activity if these spines are well constructed and functionally active (e.g. endowed with glutamatergic AMPA and/or NMDA receptor). However, surprisingly, the analysis of cumulative probability of spontaneous synaptic activity revealed a significant decrease in both the frequency and the amplitude of the spontaneous excitatory postsynaptic currents (sEPSC, Fig. 4 b, c) in L2/3 pyramidal neurons of biopsies containing Aβ compared to no-pathology containing control. This situation was reversed in biopsies containing Aβ and tau, where an increase in the frequency of the sEPSC (by left shift of cumulative probability curve) was observed compared to biopsies with no-pathology and Aβ only (Fig. 4 b). However, the cumulative probability of sEPSC amplitude in Aβ and tau cases also remained decreased compared to controls (Fig. 4 c). Very interestingly, we found an increase in the cumulative probability of amplitude for the spontaneous inhibitory postsynaptic currents (sIPSC, Fig. 4 d) of L2/3 pyramidal neurons in biopsies with Aβ or Aβ and tau.

**Fig. 4.**
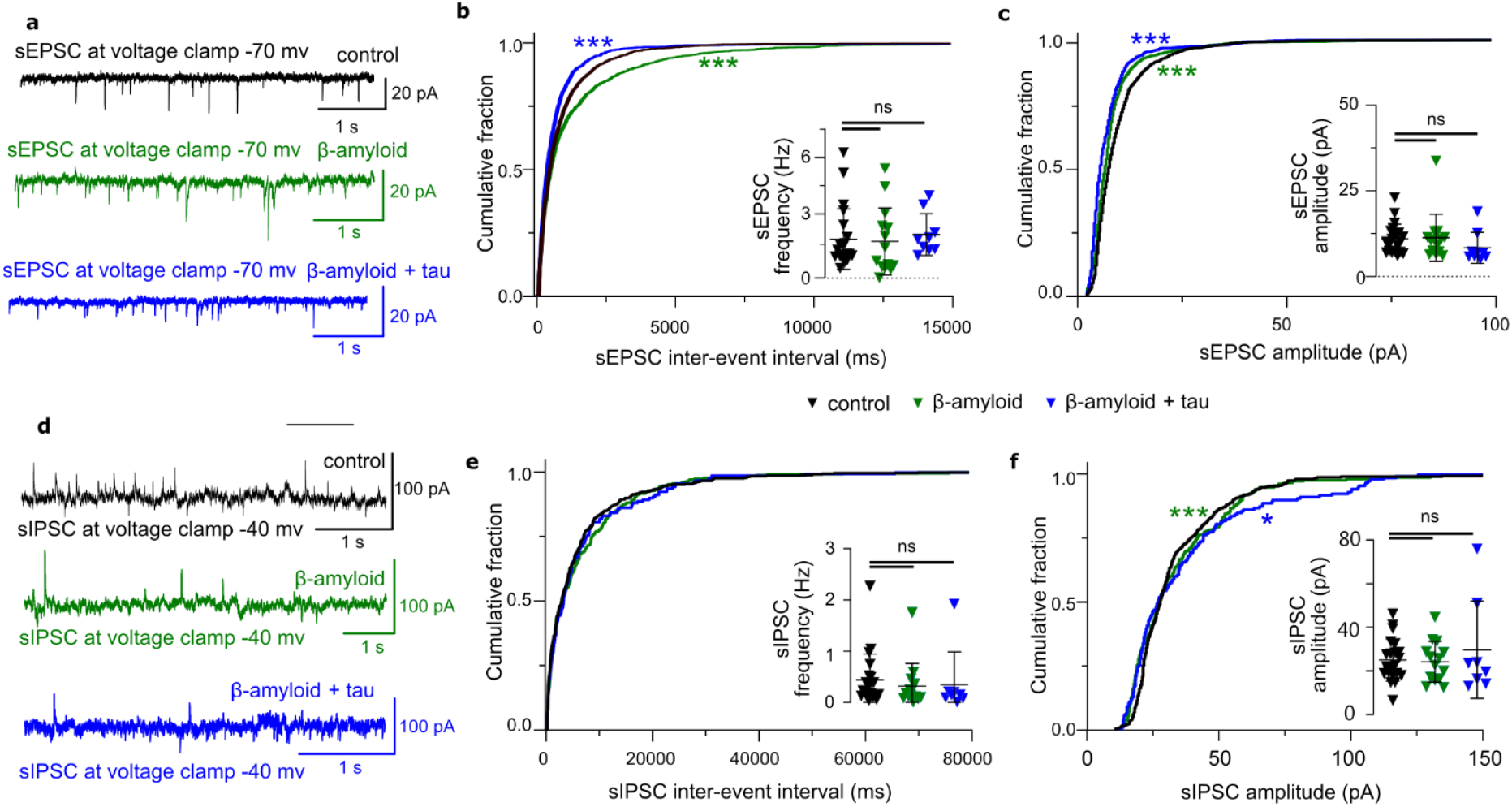
Frequency and amplitude of spontaneous postsynaptic currents in pyramidal neurons of L2/3 in iNPH patient frontal cortex biopsies. **a**. Spontaneous excitatory postsynaptic currents (sEPSC) were recorded in a voltage-clamped mode at -70 mV in different pathology cases. **b**. Cumulative probability plots of sEPSC frequency, measured by the inter-event intervals. Insets display values of the sEPSC frequency of PN in iNPH samples without Aβ (black), with Aβ (green), and with Aβ plus tau peptide (blue); **c**. Cumulative probability plots of sEPSC amplitude in correlation with Aβ pathology. Insets display box plots of the actual sEPSC amplitude of PN in iNPH patient frontal cortex biopsies. **d**. Spontaneous inhibitory postsynaptic currents (sIPSC) were recorded in a voltage-clamped mode at -40 mV in different pathological cases. **e**. Cumulative probability plots of sIPSC frequency, measured by the inter-event intervals. Insets display values of the sIPSC frequency of PN in iNPH samples without Aβ (black), with Aβ (green), and with Aβ plus tau peptide (blue); **f**. Cumulative probability plots of sIPSC amplitude in correlation with Aβ pathology. Insets display box plots of the actual sIPSC amplitude of PN in iNPH patient frontal cortex biopsies. For b, c, e and f data are presented as mean±SD. Statistical significance (*p<0.05, ***p<0.001, ns p>0.05) was evaluated by Kolmogorov-Smirnov test with Wasserstein distribution (cumulative probability plots) and Kruskal-Wallis test with Dunn’s correction (box plots), control group n=19 cells/11 biopsies, Aβ positive group n=14 cells/11 biopsies, Aβ positive plus tau group n=9 cells/7 biopsies.

### Hyperexcitability of the supra-granular layers in early AD-related pathology

To evaluate the level of excitation under different pathological conditions, we used multielectrode recording techniques (MEA) in conjunction with pharmacology as described previously^20^ (Fig. 5 a-c). We bath applied 200 μM NMDA (a glutamate receptor agonist, Fig. 5 g) over an extended period (2 minutes) and evaluated the firing responses in the cortex in a layer specific manner (Fig. 5 d). Despite the large difference of the levels of Aβ staining in iNPH biopsies (Fig.5 e), we found that there were no differences in the number of active channels responding to NMDA application amongst different patient groups (Fig. 5 f). In iNPH biopsies with Aβ only, NMDA stimulation caused a significantly larger increase in spike firing within the supra-granular layers (but not deep layers, Fig.5 h) compared to no-pathology controls. However, in biopsies containing both Aβ and tau, the response was also increased but less pronounced and statistically insignificant (Fig.5 h). Similarly, burst-like activity of sequences of spikes were observed in the cortex (Fig. 5 i) at critical frequencies (CF) above 100 Hz. Such events are known to be required for the attainment of spike backpropagation and persistent dendritic calcium oscillations in the cortex and can be indicative of the engaging and processing propensity of the cortical circuit^30–32^. We quantified the instantaneous interspike frequency and found that recordings from slices with Aβ but also in Aβ and tau, exhibited a larger proportion of ISI above CF in the supragranular layers only (Fig. 5 j-k). These results confirm that both AD-related pathology is associated with hyperexcitable cortical network states in the supragranular layers (Fig. 5 l-m).

**Fig. 5.**
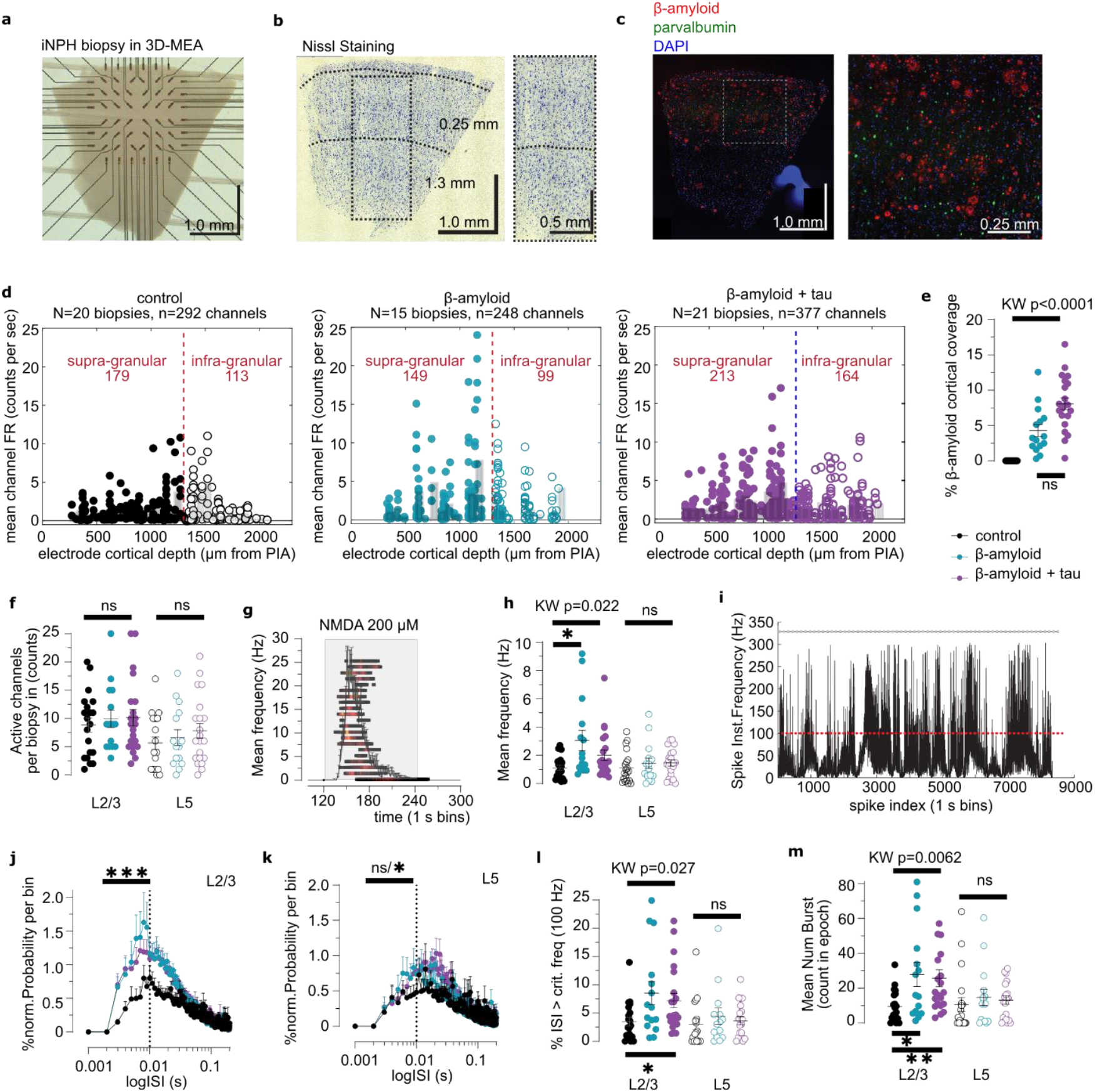
Layer specific 3D-MEA recordings from iNPH cortical slices. **a**. Image of an iNPH biopsy recorded with an 8×8 configuration 3D MEA with 60 titanium nitride (TiN) electrodes. Recording electrodes had a spacing of 0.25 mm and a height of 0.1 mm. Only the top 0.02 mm of the electrode were conductive to recording electrical signals from the intima of the slice. **b**. Left: Nissl staining of the recorded biopsy shown in **a**. Dotted lines show approximate location of L1 to L2/3 boundary and of L2/3 to L4 boundary. Right: Detail of the dotted area on the left Nissl image showing abundant pyramidal cells in L2/3 (0.5 to 1.25 mm) and sparse pyramidal cells in L5 (1.5-2.0 mm). **c**. Left: Immunohistochemical (IHC) detection of Aβ plaques & parvalbumin interneurons in the recorded slice shown in **a**. Right: Detail from IHC showing the structure of dense and diffuse Aβ plaques. **d**. Left to right: For every pathological group a plot is presented of individual channel mean firing rate (FR, NMDA-induced) as a function of cortical depth of channel location. **e**. Comparison of the % Aβ antibody coverage of the cortical area for differently grouped iNPH biopsies. Aβ positive biopsies were rank sorted according to their Aβ % coverage (burden) from histological IHC determinations to ascending values and the set was divided into two equal blocks (control negative 20 biopsies, Aβ 15 biopsies and Aβ with tau 21 biopsies). There is no significant difference between, Aβ only and, Aβ with tau biopsies in % Aβ cortical coverage (KW stat=43.47, p>0.0001, followed by pairwise Tukey’s, p>0.05). **f**. Comparison of activated channels induced by NMDA superfusion in iNPH biopsies from Aβ negative, Aβ only and, Aβ with tau burden groups. There were no significant differences in the NMDA activated channels per biopsy in either supra (0.25 to 1.3 mm cortical depth) or infra-granular layers (>1.3 mm cortical depth) (KW stat= 0.27, non-significant, ns p>0.05). **g**. Example of stacked spike raster plots overlaid on a mean firing rate plot for the iNPH biopsy exposed to NMDA shown in **a**. Color scale in spike raster encodes for high frequency firing (>100Hz). **h**. Comparison of mean L2/3 and L5 firing rates during NMDA superfusion for iNPH biopsies from different pathological groups. Significance for multigroup non-parametric statistical testing (KW) denoted with black bars above (KW stat = 7.67, p<0.05). Pairwise statistical comparisons for significant KW results between selected significantly different groups is denoted by black bars below (Tukey’s pairwise corrected multiple comparisons test, p<0.05). For inter layer comparisons within groups, Aβ biopsies, but not any other groups, had significant higher FR in L2/3 than L5 channels (Wilcoxon non-parametric paired test, W sum of signed ranks =-82, p<0.05). **i**. Spike firing instantaneous frequency plot from concatenated spike intervals of all active electrodes from the biopsy shown in **a**. The red line represents a cut off critical frequency of 100 Hz, grey line represents the dead time interval (0.0033 s, 333 Hz) for spike detection. **j**. Histogram of normalized probability per bin against inter-spike interval (ISI) between different pathological groups for L2/3 spiking electrodes per biopsy. ISIs above a critical interval (0.01s== 100Hz) are depicted with the vertical black line. Asterisks denote strong significance in the likelihood of occurrence of low (high frequency) ISI intervals between the negative and the Aβ groups (two-way ANOVA, comparison of logISI bins, F _(998, 52947)_ = 40.41, p<0.0001, followed by Tukey’s corrected multiple comparison test, **p <0.01, ***p <0.001). **k**. As above in **e** but for L5 residing electrodes per biopsy (two-way ANOVA, comparison of logISI bins, F _(998, 52947)_ = 19.28, p<0.0001, followed by Tukey’s corrected multiple comparison test, *p<0.05, ns p>0.05). **l**. Comparison of the percentage occurrence of ISIs above critical frequency per biopsy for different pathological groups (KW stat = 7.21, *p<0.05). Pairwise significance between groups denoted with a black bar below the relevant groups (Tukey’s pairwise corrected multiple comparisons test, *p<0.05). For inter layer comparisons within groups, Aβ with tau biopsies but not any other groups, had significant higher occurrences of ISIs with larger than critical frequency in L2/3 than L5 channels (Wilcoxon non-parametric paired test, W sum of signed ranks =-128, p<0.05). **m**. Comparison of the bursting rate (count in drug epoch) per biopsy for different pathological groups. Burst start was detected when the ISI between two spikes fell below 0.01 s (>100 Hz). and terminated when the intra-burst frequency fell above 0.02 s (<50 Hz). Bursts with less than 0.025 s inter-burst interval were merged (KW stat = 10.17, **p<0.01, Tukey’s pairwise corrected multiple comparisons test, **p<0.01). For inter layer comparisons within groups, Aβ with tau biopsies but not any other groups, had significant higher BR in L2/3 than L5 channels (Wilcoxon non-parametric paired test, W sum of signed ranks =-106, *p<0.05).

## Discussion

The electrophysiological properties of pyramidal neurons in the L2/3 cortical layer from biopsies of patients with iNPH containing Aβ and tau provide compelling insights into the early pathophysiological alterations observed in AD. Our findings reveal that the presence of Aβ and tau distinctly modulate the electrical properties and network dynamics of L2/3 pyramidal neurons, with significant implications for synaptic activity, membrane excitability, and neuronal communication. Our study revealed that progressive changes occur in intrinsic ionic properties, synaptic transmission and spine density in supragranular pyramidal neurons.

We failed to detect differences in the ionic properties of pyramidal neurons in biopsies with low to moderate cortical Aβ burden compared to no-pathology cases. In contrast, changes in active and passive membrane properties were observed only in biopsies with co-occuring tau pathology as shown for voltage-gated Na+ and K+ currents and membrane capacitance. The reduction in L1 induced inhibition in the supragranular layers in cases with Aβ pathology is in good agreement with the lack of functional interneuron input hypothesis proposed in earlier study^20,21^. Decreasing inhibitory inputs from L1 interneurons are expected to give rise to hyperexcitability in the supragranular tissue in early AD. We confirmed this hypothesis using MEA recordings where both spike firing frequency but also the bursting propensity above critical frequencies (representing the potential for engagement of the cortical network^31,32^) were increased in biopsies with AD-related pathology. Interestingly, the deficits in L1-induced inhibitory drive, local excitatory input (frequency) and supragranular firing hyperexcitability detected in cases with Aβ pathology only are reversed in cases with both Aβ and tau suggesting that the presence of tau may lead at least partially to restorative changes. It is hard however to safely dissect out the contribution of tau against the background effect of variable Aβ burden.

Our morphological results revealed an increase in spine density in supragranular layer pyramidal neurons in cases with AD-related pathology. This increase was primarily localized to the apical dendrites, which are responsible for receiving inputs from multiple cortical layers. Although, our NMDA activation data show clearly that the supragranular network in cortical slices is hyperexcitable in AD-related pathology, we cannot tell from our experiments to what extend have the newly formed spines contributed to this effect. More experiments are required to assess whether this early response to Aβ burden triggers functional spines, establish their receptor content and study their properties.

The increase in global spine density and the bidirectional changes in the frequency and amplitude of the excitatory transmission in cases with AD-related pathology are reminiscent of coordinated changes often observed in homeostatic plasticity and in particular in synaptic scaling^33^. In this context, an early presynaptic dysfunction caused by accumulating amyloid fragments can be seen as the initial step in a cycle of compensatory changes in synaptic signaling and spine formation leading to abnormal cortical hyperexcitability as described in early AD related pathology. Such events are expected to have a close cross talk with astrocytic and microglial microenvironment that respond to the local inflammation cause by Aβ fragments. The progression of pathology to late AD seems more involved and the potential ‘restorative’ benefits of emerging tau pathology in the presence of Aβ pathology as seen by us here (L1 inhibition, presynaptic excitation, supragranular firing) and others^20,21^ (see L1 NDNF neurons^20^) are interesting features of the overall transformation of the cellular and network cortical signatures with progressing Aβ pathology before ensuing cellular degeneration and death.

## Methods

### Primary biopsy cohort

The cohort used for patch-clamp analysis included biopsy specimens from 43 patients who were evaluated for adult hydrocephalus with NPH symptoms at the Kuopio University Hospital. The 43 individuals with idiopathic normal pressure hydrocephalus included 24 males and 19 females with a mean age of 73 years (standard deviation: 7 years), all of Finnish ancestry. Assessment of biopsy samples by a neuropathologist under light microscopy identified 15 with Aβ plaques (Aβ+), 10 with both Aβ plaques and phosphorylated tau pathology (Aβ+Tau+), and 18 biopsies that had neither histopathology. Patients consented to retrieval of brain biopsies during ventriculoperitoneal shunt placement for treatment of their symptomatic adult hydrocephalus. The biopsy procedure was approved by the Research Ethics Committee of the Northern Savo Hospital District (decision No. 276/13.02.00/2016).

### Procurement of frontal cortex brain biopsies

Biopsies were taken at the site where the shunt would penetrate the brain. A pyramid shape cortical biopsy was extracted by four sharp cuts of a knife and gentle lift by dissector to the most superficial brain parenchyma. Each cut is around 3.0 mm wide, 1.5 mm from the centre of the aimed ventricular catheter insertion site, 3.0 mm in depth, 3.4 mm in length and inclined around 65° from the surface of the brain. The insertion point of the catheter was approximately 3 cm from the midline and anterior to the coronal suture ^34^. Biopsy Ab plaques burden was further assessed semiquantitatively by a neuropathologist (Tuomas Rauramaa) under light microscopy and assigned to mild (1), moderate (2), or severe amyloid burden (3) as described previously^35^. Our initial cohort included 44 individuals. Neuroanatomical localization of biopsy was performed as described earlier^20^.

### Brain slice preparation for electrophysiological experiments

Brain biopsies were collected from the right frontal lobe from a standard insertion site (mid pupillary line, 2mm anterior to the coronal suture representing Brodmann area 8) before insertion of the intraventricular catheter for cerebrospinal fluid (CSF) shunt as described previously^5,34^. A pyramid/rectangular shaped right lobe cortical biopsy of around 3 mm in edge was extracted by sharp cut of a knife and gently lifted and placed into a sterile 50mL falcon tube containing ice-cold, sterile filtered, N-methyl-D-glucamine (NMDG)-based artificial CSF solution (aCSF; see below for composition)^36^. The brain biopsies were brought to the neurophysiology laboratory within 15 to 20 minutes of excision. Brain biopsies were gently cleaned of any debris or blood clots by careful trituration with a large bore fire polished glass Pasteur pipette, photographed for sample size documentation and were then embedded in 2% TopVision low melting point agarose (Thermo Fisher Scientific Baltics, Lithuania) and sliced (350 mm) using a vibratome (Campden Instruments, model 7000 smz) in chilled (2-4 °C), fully carbogenated (95%/5%, O_2_/CO_2_) aCSF of the following composition (in mM): 92 NMDG-methyl-D-Glucamine, 2.5 KCl, 20 HEPES, 25 NaHCO_3_, 1.25 NaH_2_PO_4_, 3 Na-Pyruvate, 2 Thiourea, 5 Na-Ascorbate, 7 MgCl_2_, 0.5 CaCl_2_, 25 Glucose (pH adjusted to 7.3 with HCl 10 M). We routinely discarded the first and last slice (extremities of the sample) obtained and only two mid-sample mini slices (total area of 6-10 mm^2^) from each biopsy were included in our study for functional examination. After cutting, slices were photographed once again and were then placed in a custom-made chamber and allowed first to recover at 34 °C for 45 minutes in the recording solution (see below for composition, supplemented with 3 mM Na-Pyruvate, 2 mM Thiourea and 5 mM Na-Ascorbate) and then for another 60 minutes in the same solution at room temperature (20-22°C) before use for electrophysiology.

### Whole-cell electrophysiology

A single slice (350 μm) was transferred and secured with a slice anchor in a large volume bath under an Olympus BX50WI microscope equipped with differential interference contrast (DIC) optics, a x40 water immersion objective and a charge-coupled device (CCD) camera (Retiga R1, Q-imaging). Slices were continuously perfused at a rate of 2.5-3 ml/min with a recording solution of the following composition (in mM): 120 NaCl, 2.5 KCl, 25 NaHCO_3_, 1.25 NaH_2_PO_4_, 2 CaCl_2_, 1 MgCl_2_, 25 Glucose. Whole-cell current and voltage-clamp recordings were conducted at 32-33 °C with an Axopatch-200B amplifier (Molecular Devices) using 5-8 MΩ glass electrodes filled with an internal solution containing (in mM) 135 Potassium-gluconate, 5 NaCl, 10 HEPES, 2 MgCl_2_, 1 EGTA, 2 Mg-ATP, 0.25 Na-GTP & 0.5% Biocytin (pH adjusted to 7.3 with osmolarity at 275-285 mOsm/l). Electrophysiological data were low pass filtered at 1 KHz (4-pole Bessel filter) then captured at 10 KHz via a Digidata 1440A A/D board to a personal computer, displayed in Clampex software (version 10.7, Molecular Devices) and stored to disk for further analysis.

### Electrophysiological protocols

Cell capacitance was measured during manual compensation on Digidata 1440A amplifier. Sodium, potassium, leak and hyperpolarization-activated cation (HCN) currents were examined under voltage-clamp through incremental voltage steps from suitable holding potentials (sodium & sustained potassium current, holding voltage of -70 mV, steps of +10 mV for 1s to +30 mV; HCN current, holding voltage of -50 mV, steps of -10 mV for 1s duration to -140 mV, Extended Data Fig. 1 a, b) as described previously ^24,27^. To examine current-input spike-output firing relationships and subthreshold currents, the neurons were given, 1s incremental current injections of ± 5-50 pA from their resting membrane potential. AP characteristics such as threshold, amplitude and afterhyperpolarization (AHP) amplitude were measured from AP threshold (defined as the point where dV/dt exceeds 10 mV/ms, Extended data Fig. 1 b) while rheobase was determined from the minimum required injected current to evoke spiking into the neuron from it resting membrane potential, measured earlier at 0 current level (Extended data Fig. 1 b, c). For every neuron, the own rheobase was taken as a starting point for AP production and as 100 % ^24^. The load survey study was in the form of current input where pulses of plus 100%, 200%, or 300% to rheobase were used to evoke firing at various frequencies (Extended Data Fig. 1 c, d).

Excitatory and inhibitory spontaneous synaptic current (EPSCs and IPSCs) were sampled at a holding potential of -70 mV and to -40 mV (to avoid recruitment of APs) respectively in 2 minute epochs as reported previously^29,37^. Synaptic stimulation for cells expressed in L1 layer was achieved using a customized stimulating electrode (round bipolar, 125 µm diameter of pole, 150 µm interpole distance, FHC Inc) positioned laterally to the L1 layer (within 200 µm from the pial surface). We delivered bipolar stimulus currents of 200-400 µs in duration and of 0.5-1-2 mA in amplitude to the slice through a constant current stimulus generator (DS-3, Digitimer). The L1 stimulation protocol was performed in current-clamp mode under application of a holding current 0 pA with step +50 pA for 7.8 s duration to +350 pA. The stimulation was applied at 1.6 s after the base activity recording and then repeated twice more with 2s interval (Fig. 2).

Cumulative probability for inter-event interval and amplitude constructed using 100 events for sEPSC and 20 events for sIPSC. The miniature amplitude threshold was set at 4 pA. The Kolmogorov-Smirnov test with Wasserstein distribution test was used to measure the statistical significance of the cumulative probability plot ^38^.

### Biopsy & neuron exclusion-inclusion criteria

iNPH brain biopsies, from which we have successfully recorded neurons, were only selected for inclusion in our database after careful consideration of experimental notes, biopsy documentation images at the time of slicing, Nissl staining & Neurolucida morphometric reconstruction records (see below) in order to ascertain the preservation of pia to L4/5 cortical layers in the biopsy slice. From such samples with well-preserved cortical layers, we accepted for electrophysiological analysis only pyramidal neurons of L2/3 that had a maximal somatic depth ranging from 450 µm up to 1200 µm from the pial surface. Somatic depth was ascertained by both biocytin-filling and IR-DIC microscopic depth calculations during recordings. Accepted pyramidal neurons of L2/3 exhibited electrophysiological stability during recordings with a stable series resistance (R_s_) of less than 30 MΩ and a stable input resistance (R_in_) with a less than 15% change after whole-cell break-in and during the recording. Accepted neurons also had to exhibit largely intact dendritic morphology devoid of aberrant or large truncations to apical-basal dendrites and this was assessed by the development and imaging of biocytin-filled recovered cells from each slice.

### DAB-staining, reconstructions, and plaque detection

Biocytin-filled neurons from recorded slices were subjected to DAB staining and neuromorphometric reconstructions as reported previously^28^. Briefly, slices were fixed in fresh 4% PFA for 48-72 hours at 4 °C and then were stored indefinitely in a PBS with 0.1% sodium azide at 4 °C until processed. Reconstructions were performed with a calibrated XYZ axis Ludl electronics stage under an x100 oil objective of a Nikon Eclipse microscope using Neurolucida software (version 10, MBF Biosciences). For reconstructions, we selected only L2/3 neurons where the apical and the basal trees were largely preserved intact. Analysis of the morphological features (e.g. total dendritic length, total nodes, spine density) for neurons was conducted via Neurolucida Explorer software.

After the neuromorphology reconstruction the slices were rewashed in Xylen and restained for primary immunopositive WO-2 antibodies (2 days overnight incubation, MABN10, Millipore, dilution 1:1000, at room temperature) and followed by a two hour incubation with an anti-mouse fluorescent secondary antibody (Alexa Fluor Goat-anti-mouse 488, A11001, Thermo Fischer Scientific, dilution 1:500). Imaging of Ab plaques situated within the biocytin-filled neurons were performed with LEICA thunder imager 3D tissue slide scanner (Cell and Tissue Imaging Unit, UEF) with minimal z interval during the stack scanning. Then the zone of contact between Ab plaque and dendrite was again rechecked in Neurolucida software for the defining of local spine density values.

### 3D-multielectrode array electrophysiology

A single slice (350 μm) was transferred and secured with a slice anchor in the chamber of a 60-3D-multielectrode recording array (3DMEA, Multichannel Systems [MCS]) with an 8×8 configuration of pyramidal electrodes bearing titanium nitride (TiN) tips (impedance 150-300 KOhm) and silicon nitride isolation. Pyramidal electrodes had a height of 100 mm but were conducting only on top 20 μm of the tip which measured 12 μm in base. Recordings were made with a MEA2100-Mini headstage (MCS, Germany). Slices were continuously perfused at a rate of 3-3.5 mL/min with a recording solution of the following composition (in mM): 120 NaCl, 2.5 KCl, 25NaHCO_3_, 1.25 NaH_2_PO_4_, 2 CaCl_2_, 1 MgCl_2_, 25 Glucose. Slices were allowed at least 30 minutes of settling time in the MEA chamber before any recordings or pharmacology was attempted. Drugs (NMDA, 200 mM for 2 minutes, Sigma-Aldrich) were bath applied by continuous perfusion via a peristaltic pump. Electrophysiological data were band passed between 1 and 3500Hz (second order Butterworth filter) and were captured at 20 KHz via MCS-IFB 3.0 multiboot (MCS, Germany) to a personal computer, displayed in MCS Experimenter software (version 2.15) and stored to disk for further analysis.

### 3D-MEA data analysis

NPH brain biopsies were selected for neurophysiological analysis after careful consideration of experimental notes, biopsy documentation images at the time of recording & Nissl staining to ascertain the preservation of pia to L4-L5 cortical layers. Multielectrode recording data files (.msrd) were imported in Neuroexplorer (Nex Technologies, US) and opened in Neuroexplorer (Nex Technologies) or MATLAB (The MathWorks) for filtering, down-sampling, measurement routines and analysis with custom scripts (python, MATLAB). Multichannel signals were analyzed for spike activity based on their cortical depth (L2/3: 300-1250 mm; L5: >=1500 mm;). Spike detection of multiunit activity (MUA) was performed from drug responding, channels of interest (COIs). Reference channel noise was subtracted from each COI before applying a 300-3500Hz band-pass filter (3rd order, Butterworth) to the raw signal followed by a spike detection threshold set at -5.5 times the standard deviation (SD) of the baseline of each COI. Electrodes qualified for analysis if they responded to NMDA drug superfusion with an increase in firing (excitation) consisting of a minimum of two consecutive 10s binned frequency peaks above 0.1Hz and a mean firing rate during NMDA application of more than 0.05 Hz. Extracted spike timestamps were stored in a file and were used to compute interspike intervals (ISI) & spike bursting characteristics for each COI during pharmacological experiments with NMDA. For burst detection, we used an interval-based algorithm to detect and quantify the characteristics of bursting induced by NMDA superfusion. We defined a cortical burst by a minimum of two spikes occurring within 10 ms of each other (minimum burst start with instantaneous doublet frequency of > 100 Hz)113,114 with burst terminating only if the forthcoming spike occurred more than 20 ms after the last (termination instantaneous frequency of < 50Hz), while any two consecutive bursts had to be separated by a minimum of 25 ms from each other (minimum interburst interval).

### Histological determinations and immunohistochemistry

A single, 350 mm, biopsy slice (3D-60MEA recorded) was fixed in fresh 4% PFA for 24-72 hours at 4 °C and then stored in PhosphateBuffer (PB) containing 30% sucrose for 24 hours at 4 °C for cryoprotection. Slices were then resliced into 20 mm sections at -20 °C using a cryostat (Leica CM1950, Leica Biosystems, Germany) and were collected on frost slides for Aβ staining. We used a standard citrate buffer antigen retrieval protocol (with 10 mM sodium citrate) prior to overnight incubation of slices with WO-2 primary antibody (MABN10, Millipore, dilution 1:1000, at room temperature) followed by a two hour incubation with an anti-mouse fluorescent secondary antibody (Alexa Fluor Goat-anti-mouse 568, A11004, Thermo Fischer Scientific, dilution 1:500) for Aβ detection. NPH biopsy slices were imaged with LEICA thunder imager 3D tissue slide scanner (Cell and Tissue Imaging Unit, UEF) and the quantification of Aβ burden was conducted in ImageJ. Regions of interest (ROIs) were drawn to outline the sample sections and Aβ burden was quantified by measuring the fluorescence intensity and area coverage fraction (intensity per mm2) of the WO2 staining inside each ROIs. Acquired images were first turned into 8-bit images, thresholded to minimum value of 30 and normalized to a maximum value of 120.

### Software and Statistical Analysis

Single-cell electrophysiological current and voltage-clamp data were analyzed in Clampfit (Molecular Devices) while synaptic data were detected and analyzed with Mini Analysis software (Synaptosoft Inc) as reported previously^29,39^. All results were exported to Prism (version 8.02, GraphPad US) for graphing, correlation, and statistical analysis. Parameter values were tested for normality of distribution through D’Agostino-Pearson or Shapiro-Wilk testing and correlated against the values of other parameters with a Spearman non-parametric test. Pairwise comparisons of values for any parameter were conducted with a two-tailed, multiple ANOVA comparisons were conducted with Kruskal-Wallis tests (followed by Dunn’s multiple comparison tests) while comparisons of cumulative distributions were evaluated with a Kolmogorov-Smirnov test with Wasserstein distribution. The comparison of spine density obtained by Neurolucida morphometric measurements was conducted with Kruskal-Wallis tests with the false discovery rate correction (Benjamini, Krieger and Yekutieli method) to prevent false positive effect, possible with manual measurements. We compared the Aβ negative samples (Aβ score 0) against a single pooled group of Aβ positive samples (for increasing the sample size) or against Aβ and tau positive sample group. Values reported herein represent mean ± standard deviation (SD). Statistical probability is inferred when P value is less than 0.05. A P value of more than 0.05 and less than 0.1 denotes a strong statistical trend in the dataset.

## Author contributions

D.A., P.A., A.P., V.L., T.M. designed the study. P.A., D.A., N.N.V., M.G.B, A.P., D.N., R.G., A.S., A.M., T.R., V.L. generated the data. P.A., D.A., N.N.V., A.P., M.G.B., M.R., L.G., R.G., T.R. analyzed the data. P.A., D.A., A.P., N.N., L.G., T.M. interpreted the data. V.L., T.M. managed the project. P.A., D.A., A.P., T.M. wrote the manuscript.

## Acknowledgments

We would like to thank Dr. Rashid Giniatullin and Dr. Jukka Jolkkonen for helpful discussions. We are also grateful for the help of Marita Parviainen and Tiina Laaksonen with patient management and cognitive testing as well as Andrea Jiang with performing the b-amyloid ELISA assays. We thank Mirka Tikkanen for technical expertise for tissue immunohistochemistry. We also thank Ylli Torn and Hanna Härkönen for acute slice physiology experiments. This work was supported by the Kuopio University Hospital VTR fund (grant 5252614 to V.L.), Academy of Finland (grants 339763 and 334801 to T.M. and 338182 and 328287 to M.H.), the Sigrid Juselius Foundation, the Strategic Neuroscience Funding of the University of Eastern Finland, the Finnish Cultural Foundation, and the North Savo Regional Fund and by the European Union (ERC, HUMANE, and 101043584 to T.M.). Views and opinions expressed are those of the author(s) only and do not necessarily reflect those of the European Union or the European Research Council Executive Agency. Neither the European Union nor the granting authority can be held responsible for them.

## Competing interests

the authors declare no competing interests.

## Requests for materials

should be addressed to Tarja Malm.

## Notes

### Competing Interest Statement

The authors have declared no competing interest.

