## Extended Data for "Early amyloid spine response and impaired synaptic transmission of pyramidal neurons in human biopsies with Alzheimer’s Disease-related pathology"

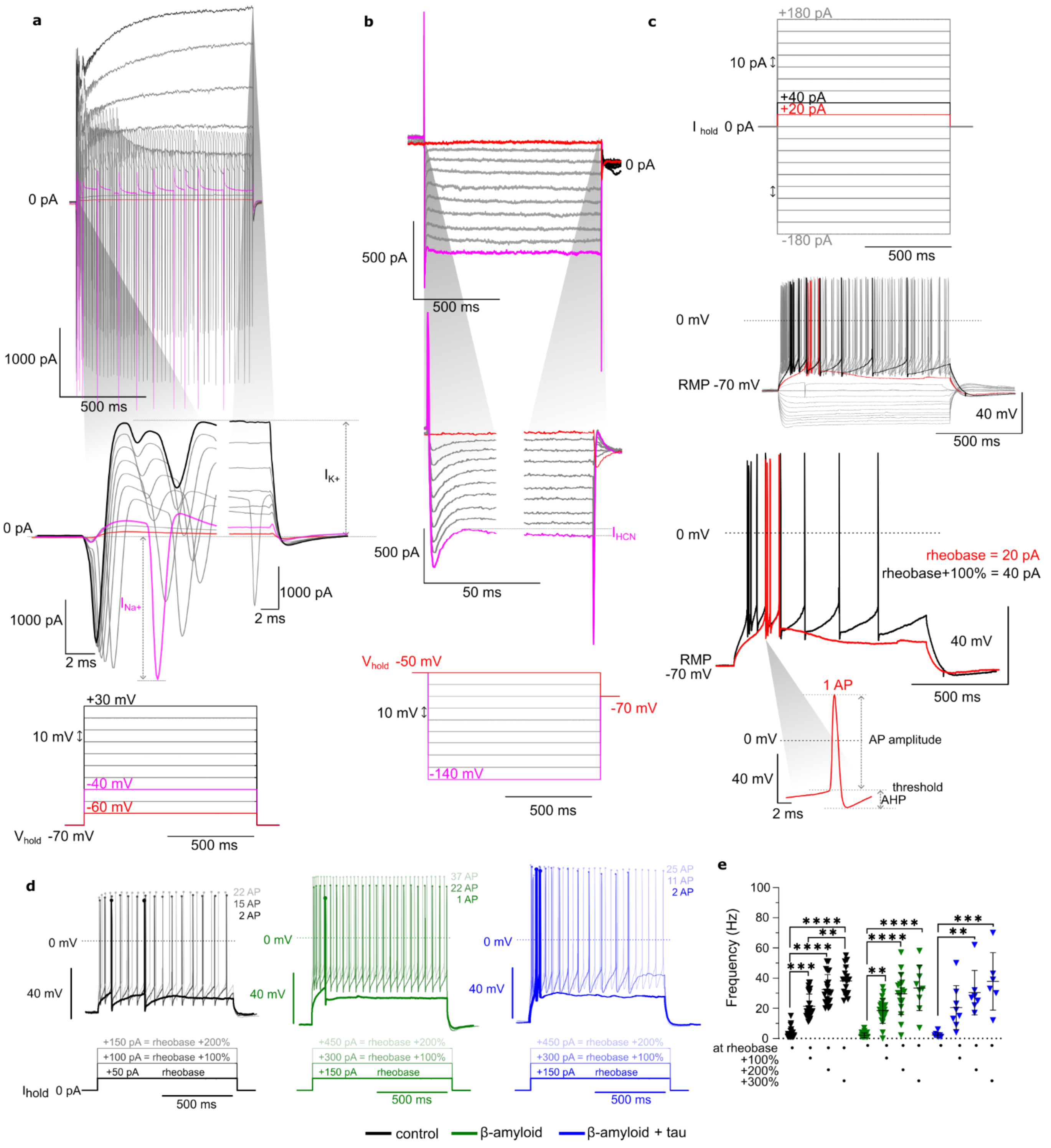


**Extended Data Fig. 1 Active electrophysiology properties. a**. Sodium ($Na^+$) and potassium ($K^+$) were examined under voltage-clamp through incremental voltage steps from suitable holding potentials: sodium & sustained potassium current, holding voltage of -70 mV, steps of +10 mV for 1s to +30 mV. **b**. Hyperpolarization-activated cation (HCN) currents were examined under voltage-clamp through incremental voltage steps from suitable holding potentials: holding voltage of -50 mV, steps of -10 mV for 1s duration to -140 mV. **c**. To examine current-input spike-output firing relationships and subthreshold currents, the neurons were given, 1s incremental current injections of ± 5-50 pA from their resting membrane potential. AP characteristics such as threshold, amplitude and afterhyperpolarization (AHP) amplitude were measured from AP threshold (defined as the point where dV/dt exceeds 10 mV/ms) while rheobase was determined from the minimum required injected current to evoke spiking into the neuron from it resting membrane potential, measured earlier at 0 current level **d**. For every neuron, the own rheobase was taken as a starting point for AP firing frequency normalisation as 100 %. The load survey study was in the form of current input where pulses of plus 100%, 200%, or 300% to rheobase were used to evoke firing at various frequencies. **e**. Frequency at the rheobase, at the +100%, +200% and +300% growth to rheobase in PN registered in iNPH samples without β-amyloid (black), with β-amyloid (green), or with β-amyloid plus tau peptide (blue). Data presented as mean±SD with statistical comparison by Kruskal-Wallis test with Dunn's correction, **p<0.01, ***p<0.001, ****p<0.0001 n=26, 20 and 8 pyramidal cells/group.

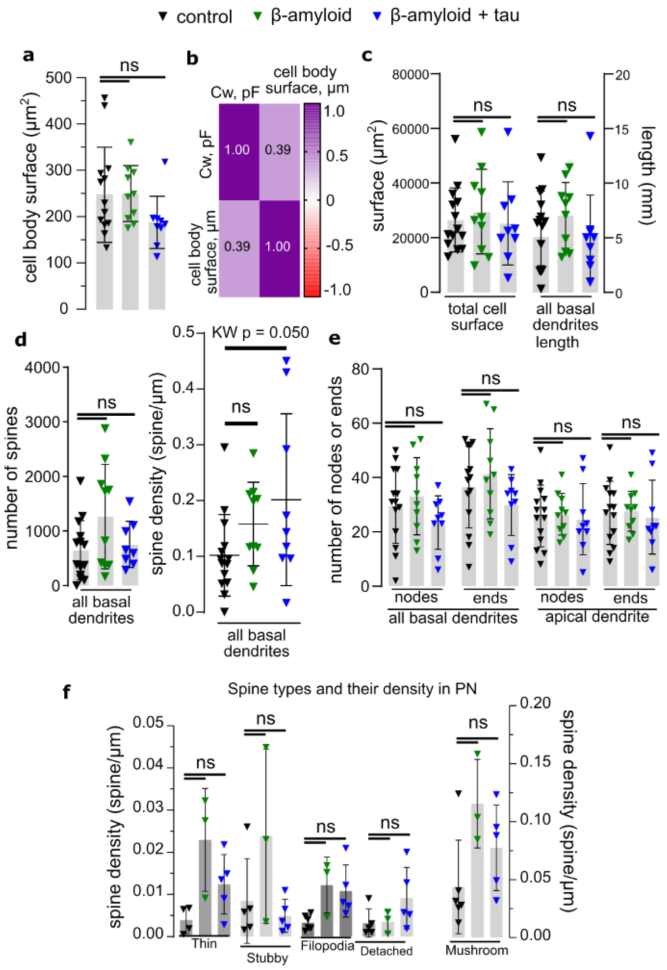


**Extended Data Fig. 2. Additional morphology reconstruction parameters of L2/3 pyramidal neurons (PN) in iNPH patient frontal cortex biopsies. a**. Cell body surface (μm²) PN registered in iNPH samples without Aβ (black), with Aβ (green) and with Aβ plus tau peptide (blue). **b.** Correlation plot of cell capacitance (pF) obtained from patched PN and their cell body surface (μm²) obtained from morphology reconstruction measurements. **c**. Total cell surface and length of all basal dendrites belonging to PN registered in iNPH samples without Aβ (black), with Aβ (green) and with Aβ plus tau peptide (blue). **d**. Number of spines and spine density detected in all basal dendrites. **e**. Number of nodes or ends of all basal dendrites or apical dendrites of PN registered in iNPH samples without Aβ (black), with Aβ (green) and with Aβ plus tau peptide (blue). **f**. Spine types and their density in PN according with Aβ presence in a tissue. For a, c-f data presented as mean±SD with statistical comparison by Kruskal-Wallis test with FDR correction, ns>0.05 and q>0.05, n=14, 10 and 9 pyramidal cells/group, taken from control negative 12 biopsies, Aβ positive 9 biopsies and Aβ with tau positive 6 biopsies.

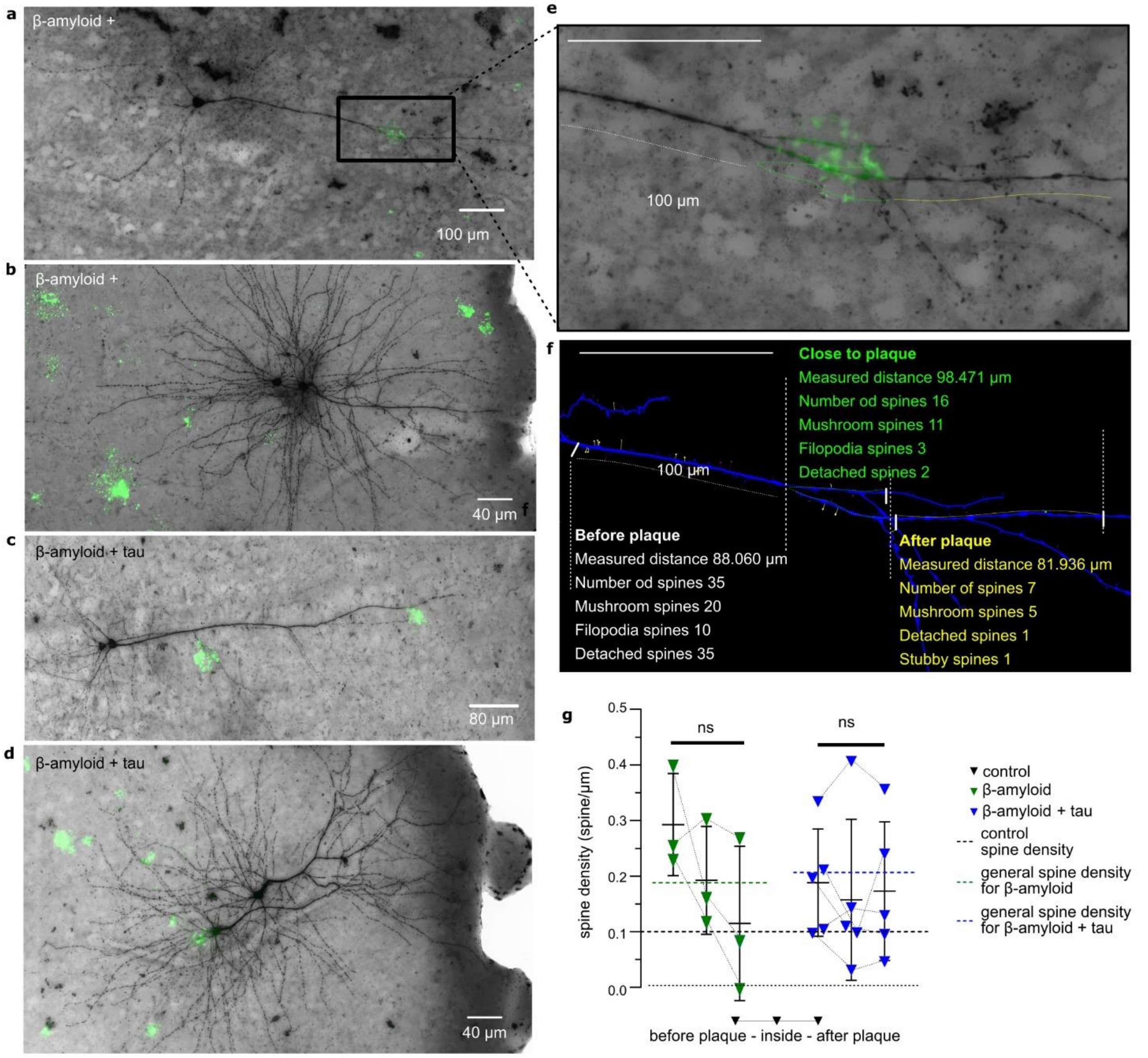


**Extended Data Fig. 3. Local Aβ plaque influence on spine density of L2/3 pyramidal neurons in iNPH patient frontal cortex biopsies a.** Aβ plaque distribution and pyramidal neurons of L2/3 in iNPH patient frontal cortex biopsies. Representative images of recorded biocytin-filled L2/3 pyramidal neurons (black) with Aβ plaques (green) marked by Alexa 488 against WO-2 monoclonal antibody in iNPH samples (**a**, **b**) with Aβ and (**c** and **d**) with Aβ plus tau peptide. Images were taken from Aβ 3 biopsies and Aβ with tau 4 biopsies. **e.** Zoomed **a** image of recorded biocytin-filled pyramidal neuron (black) with Aβ plaques (green) marked by Alexa 488 against WO-2 monoclonal antibody. **f.** Representative reconstructed trace of dendrite with spine distribution before Aβ plaque (white), close to plaque (green), and after plaque (yellow). **g.** Local spine density as before-after plot in correlation with Aβ presence in a tissue. Data presented as mean±SD with statistical comparison by Kruskal-Wallis test with FDR correction, ns $p>0.05$ and $q>0.05$, n=3 and 5 pyramidal cells/group, were 4 and 12 plaques were analyzed, taken from Aβ 3 biopsies and Aβ with tau 4 biopsies.
